# Advances and applications of the closest-tree algorithm and Hadamard conjugation in phylogenetic inference

**DOI:** 10.1101/2024.12.06.627223

**Authors:** Ernesto Álvarez González, Ricardo Balám-Narváez, Diego F. Angulo, Pablo Duchen

## Abstract

In phylogenetic inference Hadamard methods and the closest-tree algorithm have been a promising alternative to likelihood-based methods. However, applications to actual biological problems have been limited so far. In the early nineties, Hendy and Penny (1993) developed the two-state closest-tree algorithm for estimating the optimal branch lengths of a phylogenetic tree, whose parameters correspond to the Cavender’s molecular evolution model (CFN). Steel et al. (1992) then developed the four-state version of this method, whose parameters correspond to the Kimura 3ST’s molecular evolution model (K3ST). In both cases, formulas for solving the optimization problems were provided. Here, we do not only contribute with proofs for these formulas, but we also adapt this methodology to the orchid genus *Lophiarella*, whose phylogenetic relationships remain unclear. With this biological application, we show the efficacy of the closest-tree algorithm coupled with Hadamard conjugation, phylogenetic invariants and edge-parameter inequalities (in Fourier coordinates) in jointly inferring the tree topology and the molecular evolution model that best explains the data. Finally, we reconcile this phylogeny with biogeographical and morphological aspects within this genus.

## 1 Introduction

In Biology, estimation of phylogenies from DNA alignments has been mostly done using likelihood-based methods (Duchen, 2021a,b). Compared to them, Hadamard methods for tree estimation have been less applied to actual biological datasets, even though the mathematics of Hadamard transforms apply elegantly to phylogeny estimation when the number of species is not too high (Felsenstein, 2004). Similar to these traditional approaches, Hadamard methods also use a multiple sequence DNA alignment as input, but instead, each site in the alignment is categorized into patterns, and the frequencies of these patterns form “spectra” (Bryant, 2009). The closest tree algorithm is then used to infer the phylogeny that provides the closest match between observed and expected spectra frequencies (Hendy and Penny, 1993; Steel et al., 1992).

Specifically, Hendy and Penny (1993) formulated the closest tree algorithm for reconstructing phylogenetic trees under the Cavender-Farris-Neyman (CFN) model (Cavender, 1978; Coons and Sullivant, 2020) by encoding the corresponding graphs through vectors (tree specters) in a metric space. Taking Poisson distributions on the tree edges, Hendy and Penny (1993) estimated the probability of state change between the endpoints of each edge. They introduced two operations, the Hadamard-exponential conjugation and the Hadamard-log conjugation. Therein, by taking the edge lengths of the tree as parameters of the model of evolution, and the relative frequencies of the observed character patterns at the leaves as input (or external) data, they provided operations for recovering the external data when the model parameters are given, and vice versa.

Hendy and Penny (1993) provided a data-fitting formula for estimating optimal values of the tree edges, but left out a proof of it. Steel et al. (1992) then introduced the ‘conjugate spectrum’, a square matrix whose non zero entries exhibit the tree lengths of a phylogenetic tree, and used it to derive a formula for data fitting (called *Approximation formulae*), but again a proof was not provided. In this work, we contribute with proofs for the data-fitting formula of Hendy and Penny (1993) (Proposition 1) and the *Approximation formulae* of Steel et al. (1992) (Corolary 3).

Given the usefulness of Hadamard methods we were surprised to realize how little it was applied to actual biological datasets. For this reason, we delved into the (problematic) phylogeny of a neotropical orchid genus called *Lophiarella*. This study system is not only relevant from a biological point of view (i.e. its morphology, biogeography, cultivation, and conservation), but it also lends itself perfectly for our purposes, since it has a limited number of species, with unclear phylogenetic relationships that can be resolved with a combination of Hadamard conjugation and the closest tree algorithm. In the present work, we did not only provide with the “conjugate expectrum” proofs explained above, but we also used these formulas to infer the phylogeny of *Lophiarella*, as well as evaluating the performance of the Approximation formulae through the edge-length spectrum to calculate edge (or branch) lengths.

## 2 Advances in the Hadamard and closest-tree method

In what follows, we use the terms phylogenetic tree, phylogeny, or tree interchangeably. We follow the biological interpretation of a phylogeny as a representation of the evolutionary relationships between species. If the phylogenetic tree is rooted, a hypothetical reconstruction of the evolutionary process starts at the root of the tree with a single nucleotide sequence of length *L*. This sequence then “evolves” along the edges of the tree according to a model of molecular evolution, e.g. JC69, K2ST, K3ST, GM (Yang, 2006). Such model will change or “mutate” the nucleotides with a given probability. As a result, at the end of the tree, each of *n* leaves will have a final nucleotide sequence, totalling *n* such sequences. The set of nucleotide sequences at the leaves is referred as a multi-species sequence alignment (or simply, an alignment). Such alignment is represented as a character matrix where each row contains DNA nucleotides (one of {*A, C, G, T*} per matrix entry) for a given species. Thus, the dimension of an alignment is *n × L*, where *L* is the number of nucleotides, and *n* the number of species (or leaves, as stated above).

In reality, we do not know the sequences in the internal vertices of the tree, but only at the leaves. Thus, the goal is to infer or reconstruct the topology of a tree using the DNA alignment as input. Several such methods have been developed (e.g. Parsimony or Likelihood-based methods) but here we will focus on the Closest-Tree Algorithm and Hadamard Conjugation.

### 2.1 The Closest Tree Algorithm

Let 𝒯 be a phylogenetic tree with *n* leaves. The columns of the corresponding *n* × *L* alignment are called character patterns. We count all of them and get the observed vector of frequencies. We can also compute the expected vector of frequencies of character patterns in terms of the parameters. The closest tree algorithm is a leastsquares fit of the observed vector of frequencies to the expected vector of frequencies.

There are two versions of the closest tree algorithm, the 2-state of Hendy and Penny (1993) and the 4-state of Steel et al. (1992). For the 2-state version, Hendy and Penny (1993) provide a formula for the fit in Proposition 1, for which we contribute with a proof. For the 4-state version, Steel et al. (1992) provide a formula for the fit in Corollary 3, for which we also contribute with a proof.

#### 2.1.1 Two-state version

Let 𝒯 be a phylogenetic tree with *n* leaves in *N* = {1, 2, …, *n*}. We associate to it a 2^*n−*1^-dimensional vector, called *tree spectrum*, whose entries, other than the first, are non-negative. Actually, the positive entries correspond to the 𝒯-branches and the first entry equals the negative sum of the rest. Hendy and Penny (1993) define the spectral set, *W* (𝒯), as the set of all possible tree spectra of 𝒯.

The spectral space, *W* (*N*), is the union of all spectral sets associated with all possible phylogenetic trees with *n* leaves.

*W* (*N*) is a subset of the hyperplane 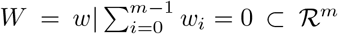, with *m* = 2^*n−*1^. Given a point *γ* ∈ *W*, we wish to find a point in *W* (*N*) closest to *γ*. The nearest tree selection procedure chooses a tree (𝒯, *w*) with *w* ∈ *W* (𝒯) that is closest to *γ*, according to the Euclidean metric. Here, we provide a proof of the following proposition presented by Hendy and Penny (1993):

**Proposition 1**. *Let γ ∈ W and 𝒯 a phylogenetic tree with leaves. Let* 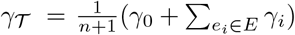, *where E is the set of 𝒯-branches. The nearest point to γ within the spectral space of 𝒯 is w* = (*w*_*i*_) *with w*_*i*_ = *γ*_*i*_ *− γ*_*T*_ *for all e*_*i*_ *∈ E (if γ*_*i*_ *> γ*_*T*_ ).

*Proof*. Let *γ*^*′*^ *∈ W* (𝒯). We wish to minimize

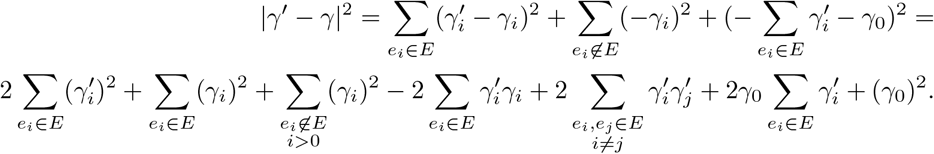

The system

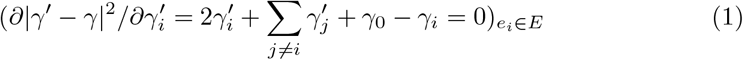

can be solved by Cramer’s Rule. This system has the matrix representation *MX* = *C*, where

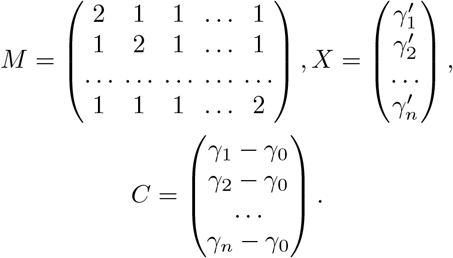

By using Equation A1 with 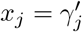 and *b*_*j*_ = *γ*_*j*_ *− γ*_0_, we get

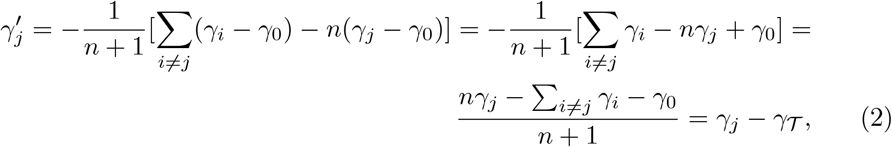

where 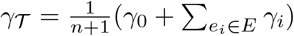.

#### 2.1.2 Four-state version

Let 𝒯 be a phylogenetic tree with *n* leaves in *N* = {1, 2, …, *n*}, whose vertices correspond to 4-state discrete random variables and whose molecular evolution model is the general model (GM).

Based on the Klein group, 𝕂 = ℤ_2_ *×ℤ*_2_, (Steel et al., 1992) define nucleotides as *A* = (0, 0), *G* = (0, 1), *C* = (1, 0), *T* = (1, 1). They identify families of character patterns (called *substitution patterns*) in terms of pairs *θ* = (*θ*_1_, *θ*_2_), whose entries are subsets of *N* ^*′*^ = *N* \{*n*}.

Let 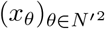 be the observed vector of frequencies for all substitution patterns in the alignment. Let 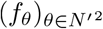 be the expected vector. Both vectors are *k* = 4^*n−*1^-dimensional. Let 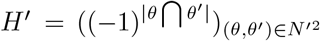 and define *H* = *H*′ ⊗ *H*′, where the operation *⊗* is the Kronecker product.

For *y* ∈ ℝ^+*k*^, let log(*y*) = (log(*y*_*i*_)) _*i*∈ [*k*]_, where [*k*] = {1, 2, …, *k*}. If (*Hx*^*t*^)_*i*_ *>* 0∀*i*, set *γ* (*x*) =*H*^*−*1^(log(*x*^*t*^)), where the super index *t* means transposition. The vector *γ*(*x*) = (*γ*_*θ*_(*x*)) is called the *conjugate spectrum*.

Substitutions are classified as transitions (*A* ↔ *G, T* ↔ *C*), type I transversions *C* ↔ *G, T* ↔ *A*) and type II transversions (*A* ↔ *C, T* ↔ *G*). Let *ρ* be an edge of 𝒯. The expected values of transitions, type I transversions, and type II transversions are denoted by 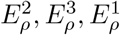, respectively.

Set 𝒞 (𝒯) = {(0, 0)} ∪ *{ρ*^(*i*)^ : *ρ ∈ 𝒯, i* = 1, 2, 3}, where *ρ*^1^ = (*ρ*, ∅), *ρ*^2^ = (∅, *ρ*), *ρ*^3^ = (*ρ, ρ*). Set *γ*^*ρ*^ = *γ*^*ρ*^ (**x**) and let 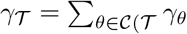.

The main theorem in Steel et al. (1992) states the following:

**Theorem 2**. *Given a distribution f* = *f* (𝒯, *P*) *for the substitution patterns on 𝒯-leaves, whose molecular evolution model is the general model, the following holds:*

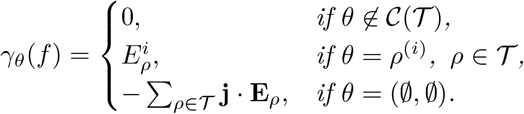

Theorem 2 uses the notation *f* (𝒯, *P*) to make precise the dependence of the distribution *f* to the phylogenetic tree 𝒯 and to the model of molecular evolution *P*. The matrix *γ*_*θ*_(*f*) of the theorem is the *edge-length spectrum* in Hendy and Snir (2008).

For the star of Figure 1, the edge-length spectrum is

**Fig. 1.**
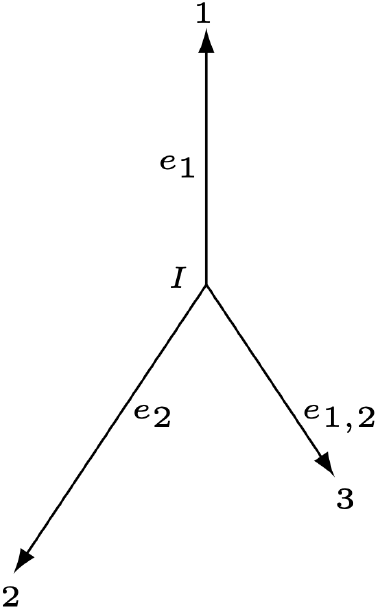
Unrooted phylogenetic tree.

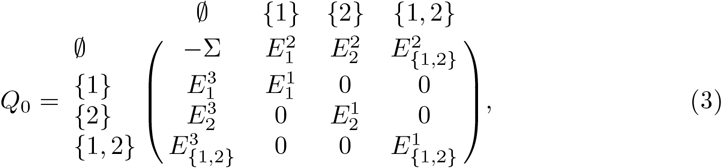

where 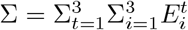.

According to Theorem 2, the entries in *Q*_0_ are expected number of substitutions.

If a phylogeny is ultrametric, that is, if it follows the molecular clock (MC) condition that it is rooted ant its leaves are aligned (or contemporary) then we denote this phylogeny as MC. For the MC comb phylogeny of Figure 2, its edge-length spectrum (with no restriction to the model of molecular evolution) is

**Fig. 2.**
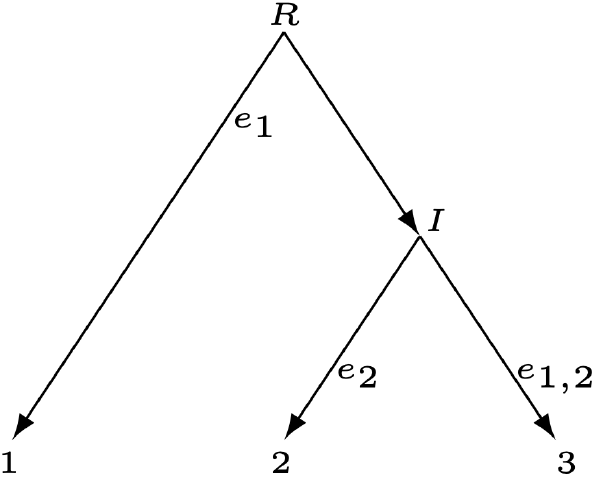
Phylogenetic tree rooted at R, where the molecular clock condition (MC) holds.

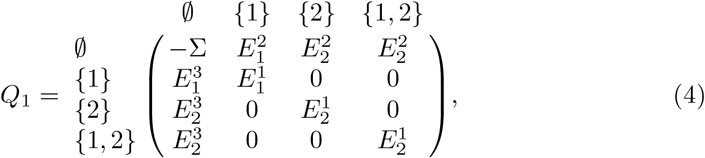

and the root is 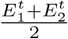 units away from any leaf of the comb for every *t ∈ {*1, 2, 3}. In this case, 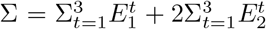. Now, we can specialize *Q*_1_ to JC69 and K2ST as follows:

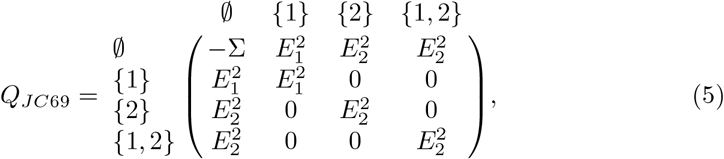

Where 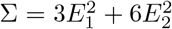, and

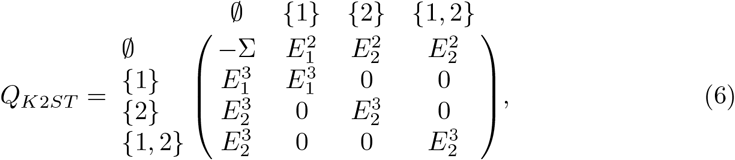

where 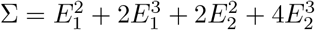.

For the tree topology 𝒯, a distribution *P* can be searched in such a way as to minimize the Euclidean distance *D*(𝒯, **x**) = ∥*γ*(*x*) −*γ*(*f* (𝒯, *P*))∥. This condition forces the parameters *E*^*i*^ to be as in Corollary 3 (see also Appendix B).

### Corolary 3

*For every phylogenetic tree 𝒯, the values* 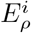 *that minimize D*(𝒯, **x**) *are given by*

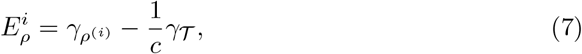

*with c* = 3|*𝒯*| + 1.

*Proof*. We wish to minimize

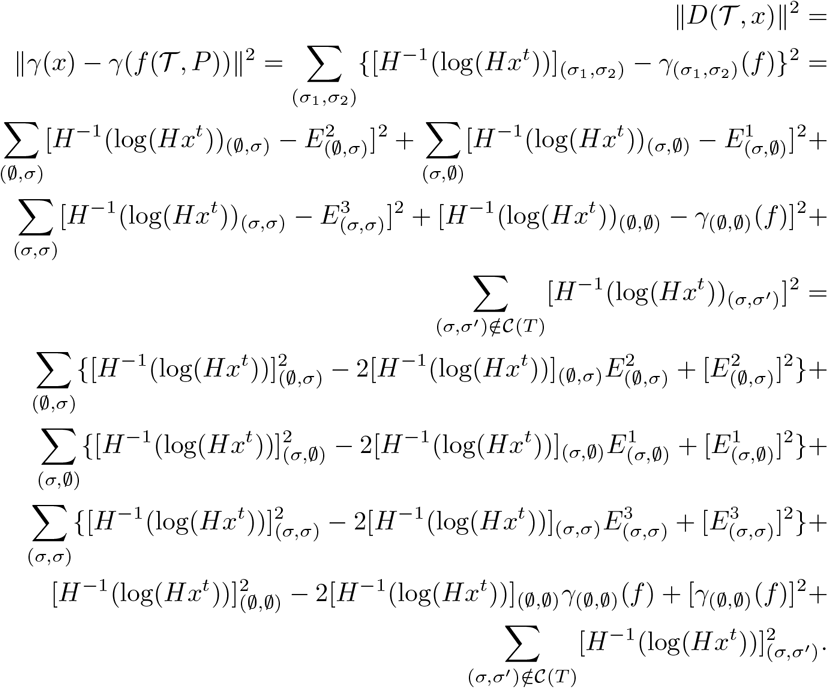

Let’s calculate the critical points of the function ∥*D*(𝒯, *x*)∥ ^2^ by partial differentiation with respect to each of the variables 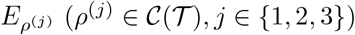 and setting them to zero. This leads to the system of linear equations

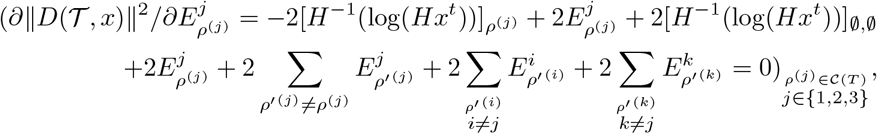

where {*i, j, k*} = {1, 2, 3}. This system has the matrix representation *MX* = *B*, with

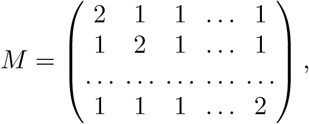

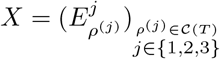, and 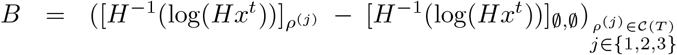, whose solution is obtained by applying Equation (A1):

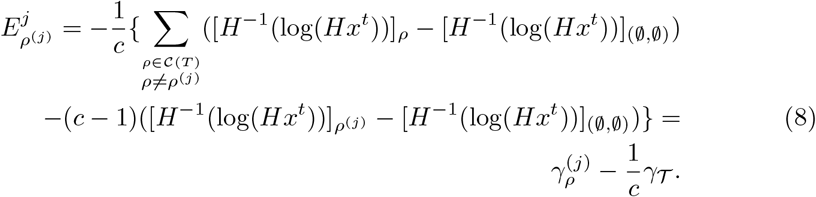

### 2.2 Hadamard Conjugation

Here, we fix the topology 𝒯 as the evolution model for the alignment of *n* species. To obtain its conjugate spectrum, *Q*, from the observed vector of frequencies, 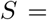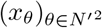 (in matrix form) as in Section 2.1.2, we use the Hadamard conjugation tool of Hendy and Snir (2008):

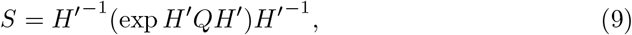

where *H*′ has been introduced in Section 2.1.2, and the exponential function is applied entry-wise.

### 2.3 Phylogenetic Invariants

Here, we use “phylogenetic invariants” to test the validity of the optimal parameter values under a model of evolution. For any given phylogenetic tree, phylogenetic invariants are polynomial functions whose variables are the probabilities of the different nucleotide sequences present in the leaves. These functions vanish on nucleotide sequences derived from the given stochastic model of molecular evolution, but are non-zero when the nucleotide sequences are derived from different stochastic models of molecular evolution Eriksson (2009).

Casanellas et al. (2005) described phylogenetic invariants in Fourier coordinates, linearly related to the Cartesian coordinates through a Hadamard matrix.

For a comb tree, let 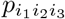 be the probability of observing the coloration *i*_1_*i*_2_*i*_3_ on its leaves, where *i*_*k*_ ∈ {*A, C, G*}, *T* for *k* = 1, 2, 3. The joint probability 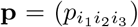 has 4^3^ coordinates. In Fourier coordinates, we have **q** = *H*′^*−*1^ ⊗ *H*′^*−*1^ ⊗ *H*′^*−*1^**p**^*t*^, where **p**^*t*^ is the transpose of **p**.

Following Casanellas et al. (2005), two probabilities 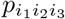 and 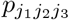 are *equivalent* if their defining polynomials are equal. Different equivalent probabilities form classes. These classes are groups of probabilities, and they are represented by *p*_*j*_, where *j* is an integer.

In Fourier coordinates, 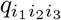 and 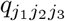 are called *equivalent* if 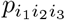 and 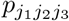 are equivalent.

Casanellas et al. (2005) provided the equivalent classes for phylogenetic trees from three to five leaves, assuming different models of molecular evolution (specializations of the K3ST model), satisfying or not the molecular clock condition (MC).

For the MC Jukes-Cantor comb tree of Figure 2 there are five classes:

Class 0 *q*_*AAC*_, *q*_*AAG*_, *q*_*AAT*_, *q*_*ACA*_, *q*_*ACG*_, *q*_*ACT*_, *q*_*AGA*_, *q*_*AGC*_, *q*_*AGT*_, *q*_*ATA*_, *q*_*ATC*_, *q*_*ATG*_, *q*_*CAA*_, *q*_*CAG*_, *q*_*CAT*_, *q*_*CCC*_, *q*_*CCG*_, *q*_*CCT*_, *q*_*CGA*_, *q*_*CGC*_, *q*_*CGG*_, *q*_*CTA*_, *q*_*CTC*_, *q*_*CTT*_, *q*_*GAA*_, *q*_*GAC*_, *q*_*GAT*_, *q*_*GCA*_, *q*_*GCC*_, *q*_*GCG*_, *q*_*GGC*_, *q*_*GGG*_, *q*_*GGT*_, *q*_*GTA*_, *q*_*GTG*_, *q*_*GTT*_, *q*_*TAA*_, *q*_*TAC*_, *q*_*TAG*_, *q*_*TCA*_, *q*_*TCC*_, *q*_*TCT*_, *q*_*TGA*_, *q*_*TGG*_, *q*_*TGT*_, *q*_*TTC*_, *q*_*TTG*_, *q*_*TTT*_

Class 1 *q*_*AAA*_

Class 2 *q*_*ACC*_, *q*_*AGG*_, *q*_*AT T*_

Class 3 *q*_*CAC*_, *q*_*CCA*_, *q*_*GAG*_, *q*_*GGA*_, *q*_*TAT*_, *q*_*TTA*_

Class 4 *q*_*CGT*_, *q*_*CTG*_, *q*_*GCT*_, *q*_*GTC*_, *q*_*TCG*_, *q*_*TGC*_

In Fourier coordinates, Casanellas et al. (2005) take the average of every representative of the given class. For example, 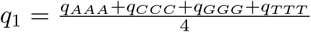.

The unique phylogenetic invariant for the MC Jukes-Cantor comb tree is

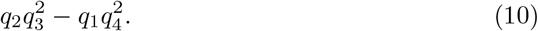

For the MC Kimura 2ST comb tree of Figure 2 there are eight classes:

Class 0 *q*_*AAC*_, *q*_*AAG*_, *q*_*AAT*_, *q*_*ACA*_, *q*_*ACG*_, *q*_*ACT*_, *q*_*AGA*_, *q*_*AGC*_, *q*_*AGT*_, *q*_*ATA*_, *q*_*ATC*_, *q*_*ATG*_, *q*_*CAA*_, *q*_*CAG*_, *q*_*CAT*_, *q*_*CCC*_, *q*_*CCG*_, *q*_*CCT*_, *q*_*CGA*_, *q*_*CGC*_, *q*_*CGG*_, *q*_*CTA*_, *q*_*CTC*_, *q*_*CTT*_, *q*_*GAA*_, *q*_*GAC*_, *q*_*GAT*_, *q*_*GCA*_, *q*_*GCC*_, *q*_*GCG*_, *q*_*GGC*_, *q*_*GGG*_, *q*_*GGT*_, *q*_*GTA*_, *q*_*GTG*_, *q*_*GTT*_, *q*_*TAA*_, *q*_*TAC*_, *q*_*TAG*_, *q*_*TCA*_, *q*_*TCC*_, *q*_*TCT*_, *q*_*TGA*_, *q*_*TGG*_, *q*_*TGT*_, *q*_*TTC*_, *q*_*TTG*_, *q*_*TTT*_.

Class 1 *q*_*AAA*_

Class 2 *q*_*ACC*_, *q*_*AT T*_

Class 3 *q*_*AGG*_

Class 4 *q*_*CAC*_, *q*_*CCA*_, *q*_*T AT*_, *q*_*T T A*_

Class 5 *q*_*CGT*_, *q*_*CT G*_, *q*_*T CG*_, *q*_*T GC*_

Class 6 *q*_*GAG*_, *q*_*GGA*_

Class 7 *q*_*GCT*_, *q*_*GT C*_

There are four invariants for the MC Kimura 2 comb tree:

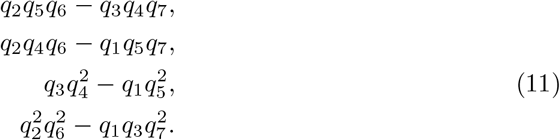

For the MC Kimura 3ST comb tree of Figure 2 there are eleven classes: Class 0 *q*_*AAC*_, *q*_*AAG*_, *q*_*AAT*_, *q*_*ACA*_, *q*_*ACG*_, *q*_*ACT*_, *q*_*AGA*_, *q*_*AGC*_, *q*_*AGT*_, *q*_*ATA*_, *q*_*ATC*_, *q*_*ATG*_, *q*_*CAA*_, *q*_*CAG*_, *q*_*CAT*_, *q*_*CCC*_, *q*_*CCG*_, *q*_*CCT*_, *q*_*CGA*_, *q*_*CGC*_, *q*_*CGG*_, *q*_*CTA*_, *q*_*CTC*_, *q*_*CTT*_, *q*_*GAA*_, *q*_*GAC*_, *q*_*GAT*_, *q*_*GCA*_, *q*_*GCC*_, *q*_*GCG*_, *q*_*GGC*_, *q*_*GGG*_, *q*_*GGT*_, *q*_*GTA*_, *q*_*GTG*_, *q*_*GTT*_, *q*_*TAA*_, *q*_*TAC*_, *q*_*TAG*_, *q*_*TCA*_, *q*_*TCC*_, *q*_*TCT*_, *q*_*TGA*_, *q*_*TGG*_, *q*_*TGT*_, *q*_*TTC*_, *q*_*TTG*_, *q*_*TTT*_.

Class 1 *q*_*AAA*_

Class 2 *q*_*ACC*_

Class 3 *q*_*AGG*_

Class 4 *q*_*AT T*_

Class 5 *q*_*CAC*_, *q*_*CCA*_

Class 6 *q*_*CGT*_, *q*_*CT G*_

Class 7 *q*_*GAG*_, *q*_*GGA*_

Class 8 *q*_*GCT*_, *q*_*GT C*_

Class 9 *q*_*T AT*_, *q*_*T T A*_

Class 10 *q*_*T CG*_, *q*_*T GC*_

There are nine invariants for the MC Kimura 3 comb tree:

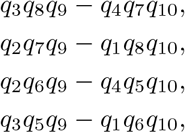

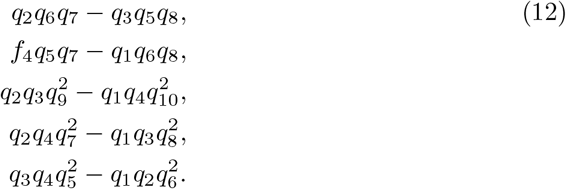

## 3 Biological application

In what follows, we will evaluate the phylogenetic invariants (Eqs. 10, 11 or 12 in **q** = *H*′^*−*1^ ⊗ *H*′^*−*1^ ⊗ *H*′^*−*1^**p**^*t*^, where **p** is the empirical distribution of character patterns from the DNA alignment of the genus *Lophiarella*. For this, we will assign its species to the leaves of Figure 2 together with each of the following molecular evolution models: JC69, K2ST or K3ST and MC. This analysis is shown in Section 3.3.

*Lophiarella* is a small monophyletic neotropical orchid. As stated above, this study system lends itself perfectly for an application of Hadamard methods because the genus *Lophiarella* has a limited number of species whose phylogenetic relationships remain unclear. The species are *L. microchila, L*.*flavovirens*, and *L. splendida*. The only existing phylogeny of this species was built using Parsimony, which resulted in a topology that is problematic with respect to morphology and biogeography (Carnevali et al., 2013) (see Discussion).

### Data input preparation

A multi-species DNA alignment was built for *Lophiarella* using the nuclear ITS gene and the chloroplast rpl32-trnL (UAG) genes. These nucleotide sequences were retrieved from the NCBI GenBank database under the following IDs: *Lophiarella flavovirens*: JQ319734, JQ366016; *Lophiarella microchila*: JQ319735, JQ366017; *Lophiarella splendida*: JQ319736, JQ366018; and the outgroup *Cohniella ascendens*: JQ366014, JQ319732. The DNA matrix (resulting from the concatenation of the above-mentioned genes) was assembled, reviewed, and edited using PhyDE v.0.99 (Müller et al., 2010). To facilitate the naming of the species, we use the notation *σ*_1_ for *Lophiarella microchila, σ*_2_ for *Lophiarella flavovirens*, and *σ*_3_ for *Lophiarella splendida*. The assembled DNA matrix has a length of *L* = 1508 nucleotides, including some gaps and some ambiguous nucleotides. The methodology we used to test the topology of the expected cladogram for *Lophiarella* is a combination of the tools: the ‘closest-tree algorithm’, ‘Hadamard conjugation’ and ‘phylogenetic invariants’. In either case, we depended on the observed ‘spectral sequence spectrum’, *S*, of *Lophiarella*, which was obtained by running the library of Álvarez (2021).

### 3.1 Implementation of the Closest Tree Algorithm

We test different tree topologies as in Figure 2 togeteher with their molecular evolution model through the application of Corollary 3. Following Álvarez and Balam-Narváez (2021), we compute the empirical spectral sequence spectra *S*_1_, *S*_2_ and *S*_3_ for the alignment orders: *σ*_1_ |*σ*_2_| *σ*_3_, *σ*_2_|*σ*_1_| *σ*_3_ and *σ*_3_ |*σ*_1_| *σ*_2_, respectively. These orders also cor002Drespond to which species is associated to what leaf of the phylogentic tree of Figure 2. For example, the alignment *σ*_3_|*σ*_1_|*σ*_2_ implies that *σ*_3_ takes the place of leaf 1 on the given phylogenetic tree; similarly, *σ*_1_ takes place of leaf 2 and *σ*_2_ takes the place of leaf 3. Matrices *S*_1_, *S*_2_ and *S*_3_ are the following

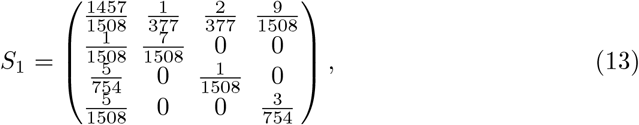

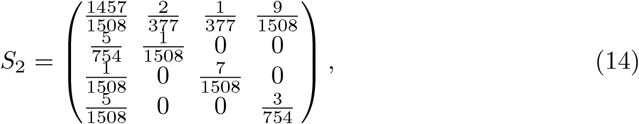

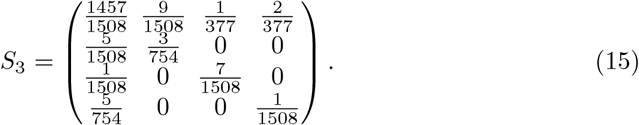

We substitute each of these matrices into Equation (9) and solve for *Q*. Then, we average the *Q*-entries according to the following assumptions:

**Assumptions for** *σ*_1_, *σ*_2_, *σ*_3_

1. The MC comb of Figure 2 as tree-topology;
2. Either model JC69, K2ST or K3ST.

For example, by setting *S* = *S*_1_ in Equation (9), we obtain

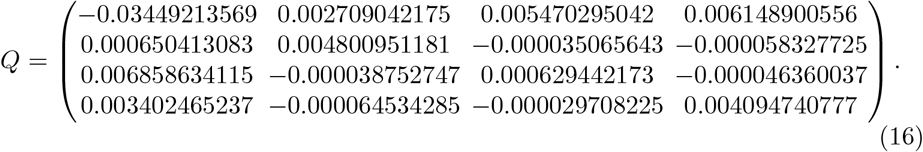

By averaging the given *Q*-entries according to the edge-length spectrum *Q*_*JC*69_ of Equation (5), we obtain

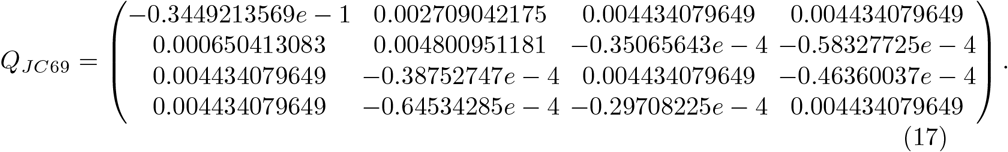

To the matrix *Q*_*JC*69_ of Equation (17), we apply Equation (7) of Corolary 3 and get 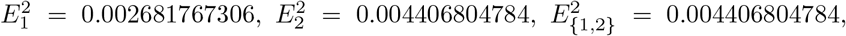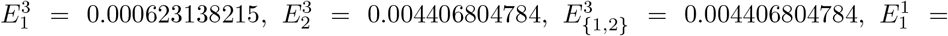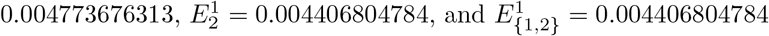.

Despite assuming the tree of Figure 2 together with (*S*_1_, *JC*69) as the model for the alignment *σ*_1_|*σ*_2_|*σ*_3_, Corollary 3 estimates transitions, type I transversions, and type II transversions along the edges. Therefore, to recover JC69 with Corollary 3 we average the terms with subscript 1, as well as the terms with subscripts 2 and 1, 2, respectively. We get 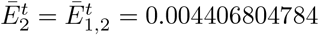 and 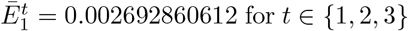 for *t* ∈ {1, 2, 3. This result contradicts the MC condition for the given alignment, since we assumed that 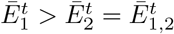. Thus, we discard the tree of Figure 2 together with (*S*_1_, *JC*69) as a plausible evolutionary model for *σ*_1_ |*σ*_2_ *σ*_3_. We proceeded similarly with the other evolutionary models present in Table 1. We emphasize that the criterion for selecting a model of evolution is that the molecular clock condition is satisfied. Table 1 shows that the evolutionary models (*S*_2_, *JC*69), (*S*_2_, *K*2*ST*), (*S*_3_, *JC*69) and (*S*_3_, *K*2*ST*) are the only models whose averages 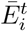 satisfy the molecular clock (MC) condition. Among these evolutionary models, (*S*_3_, *K*2*ST*) explains best the evolutionary history of *Lophiarella* (Figure 3). All of these computations are available in Álvarez (2024).

**Table 1.** Expected *Q*-entries corresponding to the empirical sequence. MEM=Molecular Evolution Model.

| S \ MEM | JC69 | K2ST | K3ST |
| --- | --- | --- | --- |
| $S_1$ | $\bar{E}_1^2 = 0.002692860612$<br>$\bar{E}_2^2 = 0.004406804784$ | $\bar{E}_1^2 = 0.002681767306$<br>$\bar{E}_2^2 = 0.005782322934$<br>$\bar{E}_1^3 = 0.002698407264$<br>$\bar{E}_2^3 = 0.003719045710$ | $\bar{E}_1^2 = 0.002681767306$<br>$\bar{E}_2^2 = 0.005782322934$<br>$\bar{E}_1^3 = 0.000623138215$<br>$\bar{E}_2^3 = 0.005103274806$<br>$\bar{E}_1^1 = 0.004773676313$<br>$\bar{E}_2^1 = 0.002334816608$ |
| $S_2$ | $\bar{E}_1^2 = 0.004292182242$<br>$\bar{E}_2^2 = 0.003607143970$ | $\bar{E}_1^2 = 0.005443020176$<br>$\bar{E}_2^2 = 0.004401696500$<br>$\bar{E}_1^3 = 0.003716763274$<br>$\bar{E}_2^3 = 0.003209867704$ | $\bar{E}_1^2 = 0.005443020176$<br>$\bar{E}_2^2 = 0.004401696500$<br>$\bar{E}_1^3 = 0.006831359245$<br>$\bar{E}_2^3 = 0.001999164292$<br>$\bar{E}_1^1 = 0.000602167305$<br>$\bar{E}_2^1 = 0.004420571112$ |
| $S_3$ | $\bar{E}_1^2 = 0.004521427322$<br>$\bar{E}_2^2 = 0.003492521430$ | $\bar{E}_1^2 = 0.006121625691$<br>$\bar{E}_2^2 = 0.004062393742$<br>$\bar{E}_1^3 = 0.003721328138$<br>$\bar{E}_2^3 = 0.003207585272$ | $\bar{E}_1^2 = 0.006121625691$<br>$\bar{E}_2^2 = 0.004062393742$<br>$\bar{E}_1^3 = 0.003375190368$<br>$\bar{E}_2^3 = 0.003727248732$<br>$\bar{E}_1^1 = 0.004067465908$<br>$\bar{E}_2^1 = 0.002687921808$ |

**Fig. 3.**
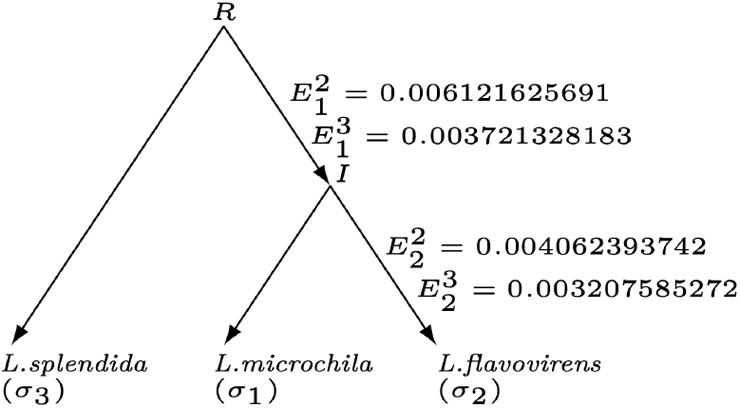
Phylogenetic Tree for *Lophiarella* under model K2ST and assuming the Molecular Clock Condition.

### 3.2 Use of Maximum Likelihood Estimation

Section 3.1 concludes that (*S*_3_, *K*2*ST*) best explains the evolutionary history of *Lophiarella* by means of the parameter values 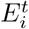 under a distance criterion. In this section, we take (*S*_3_, *K*2*ST*) as the evolutionary history of *Lophiarella*, and compute the parameter values 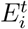 under a probability criterion. The goal is to compare both sets of parameter values. To settle this goal, we proceed analogously as Benny et al. (2006) do.

Specifically, we take *Q* = *Q*_*K*2*ST*_ in Equation (9) and solve for *S* (*Q*_*K*2*ST*_ is the matrix of Equation (6)). This is a matrix equation in the *path set variables* 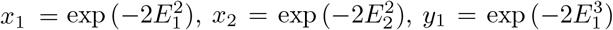 and 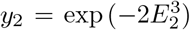 as in Benny et al. (2006). We distinguish in it exactly seven different terms: *S*[1, 1], *S*[1, 2], *S*[1, 3], *S*[2, 1], *S*[2, 3], *S*[3, 1] and *S*[3, 2].

For instance, *S*[1, 3] = *S*[1, 4], *S*[2, 1] = *S*[2, 2], *S*[2, 3] = *S*[2, 4], *S*[3, 1] = *S*[3, 3] = *S*[4, 1] = *S*[4, 4] and *S*[3, 2] = *S*[3, 4] = *S*[4, 2] = *S*[4, 3], where

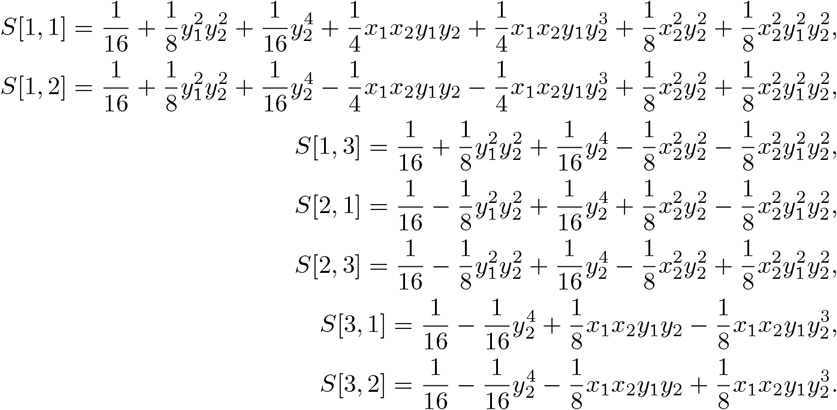

We define the likelihood function *L*(*T*) as a product of powers, whose bases are the *S*-entries and the exponents are sums of *S*_3_-entries. For example, two of the powers are 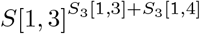 and 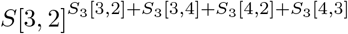. The full expression of *L*(*T*) is shown in Álvarez (2024).

We take the partial derivatives of the log version of *L*(*T*) with respect to each variable and calculate the critical points. We obtain exactly one solution for the *path set variables* with values in the interval (0, 1): *x*_1_ = 0.9879368024, *x*_2_ = 0.9919790121, *y*_1_ = 0.9926516277, and *y*_2_ = 0.9936597597 corresponding to the param-eter values 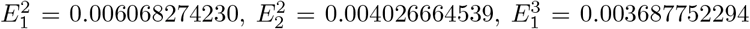 and 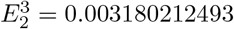, respectively. This solution satisfies the MC condition.

For these parameter values, we perform a goodness of fit test (scaled to 1508, which is the length of the alignment) as in Cochran (1952) with a significance level of 5%:

We define the classes *γ*^1^ = 1508(*S*[1, 1]), *γ*^2^ = 1508(*S*[1, 2]), *γ*^3^ = 1508(*S*[1, 3] + *S*[1, 4]), *γ*^4^ = 1508(*S*[2, 1] + *S*[2, 2]), *γ*^5^ = 1508(*S*[2, 3] + *S*[2, 4]), *γ*^6^ = 1508(*S*[3, 1] + *S*[3, 3] + *S*[4, 1] + *S*[4, 4]) and *γ*^7^ = 1508(*S*[3, 2] + *S*[3, 4] + *S*[4, 2] + *S*[4, 3]).

We compute the corresponding observations as 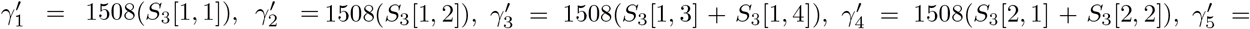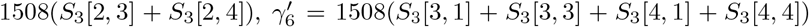 and 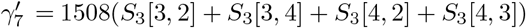.

We define the statistic

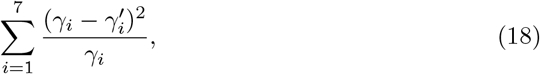

for which a realization refers to an evaluation at a point (*x*_1_, *x*_2_, *y*_1_, *y*_2_). Note that the statistic in equation (18) is class-dependent, but *S*_3_ is fixed.

The realization of the given statistic at the point (*x*_1_, *x*_2_, *y*_1_, *y*_2_), where *x*_1_ = 0.9879368024, *x*_2_ = 0.9919790121, *y*_1_ = 0.9926516277, and *y*_2_ = 0.9936597597 is 0.3862350449.

The hypothesis test is *χ*-square-distributed with 6 degrees of freedom, whose null hypothesis is 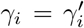 for *i* = 1, 2, …, 7. The alternative hypothesis states that, for at least one class *γ*_*i*_, *i* = 1, 2, …, 7 it holds that 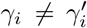. The critical value for a significance level of 5% is 12.592. Then, we do not reject the null hypothesis.

### 3.3 Use of Phylogenetic Invariants

Section 2.3 introduces Fourier coordinates, that Casanellas et al. (2005) use to compute phylogenetic invariants for small trees up to five leaves. Here, we use the phylogenetic invariants of equation (11) to confirm that K2ST best explains *σ*_3_|*σ*_1_|*σ*_2_:

The frequencies for the representative eight classes *q*_0_ through *q*_7_ are 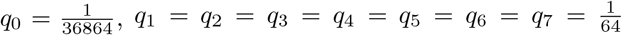. The evaluation of every polynomial in Equation (11) is zero.

According to Matsen (2009), the phylogenetic invariants for (*S*_3_, *K*2*ST*), and a set of edge-parameter inequalities must be satisfied (in Fourier coordinates) to make sure the tree edges of the tree in Figure 2 are non-negative:

We define the discrete Fourier transform for the |𝕂|-vector *f* ^(*e*)^ that estimates the transition probabilities on the edge *e* as

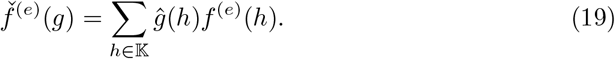

Remember that 𝕂= ℤ_2_ *×*ℤ_2_ is the Klein group as in Section 2.1.

Matsen (2009) defines the function 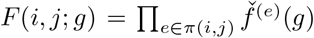, where *π*(*i, j*) is the set of edges connecting the leafs *i* and *j*, and *g* ∈ 𝕂.

A main result in Matsen (2009) is the following:

**Proposition 4**. *Given some pendant edge e, let i denote the leaf on e, and let ν be the internal node on e. Pick j and k any leaves distinct from i such that the path π*(*j, k*) *contains ν. Let w*(*g*_*i*_, *g*_*j*_, *g*_*k*_) *∈* 𝕂 ^ℒ^ *(set of edges), assign state g*_*x*_ *to leaf x for x ∈ {i, j, k} and the identity to all other leaves. Then*,

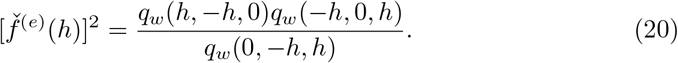

Matsen (2009) also provides the following inequalities (under model K3ST):

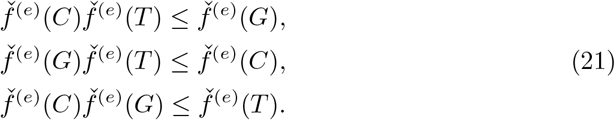

If we compute the discrete Fourier transforms as in Equation (20), then the inequalities of Equation (21) are expressed in *q*_*g*_ terms, where **q** is defined in Section 2.3. The main conclusion of Matsen (2009) is described in the following theorem:

**Theorem 5**. *If* **q** *satisfies a complete set of phylogenetic invariants (as in Section 2*.*3) for a tree T and a set of inequalities as those of Equation (21) (expressed in q*_*g*_ *terms) for each edge e* ∈ ℒ, *then* **q** *is the transform of an expected site-pattern frequency vector of T (under model K3ST) for some assignment of non-negative edge parameters to T*. *Conversely, any tree with non-negative edge parameters will satisfy such a set of inequalities*.

**Corolary 6**. *It can be easily verified that the inequalities of Equation (21) (under model K2ST) hold without any change. Then, Theorem 5 also holds in this case*.

**Corolary 7**. *According to Matsen (2009)*, (*S*_3_, *K*2*ST*) *(together with the phylogenetic tree of Figure 2) explains Lophiarella*.

*Proof*. We produce inequalities as in Equation (21) for each pendant leaf *i* as in Proposition 4 for *Lophiarella*:

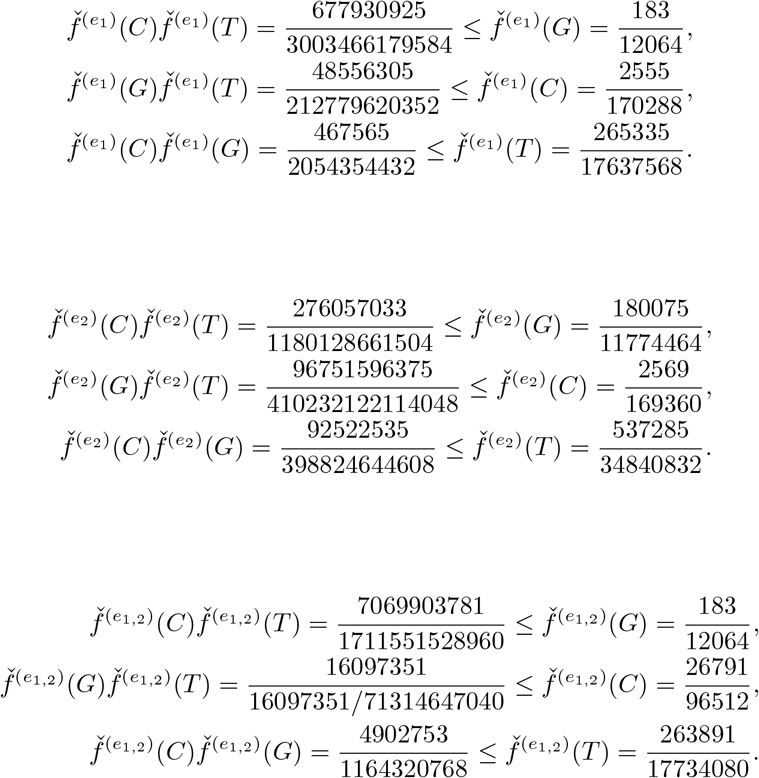

## 4 Discussion

The present work lays on the hypothesis that two of the species *σ*_1_, *σ*_2_, *σ*_3_ are more closely related. We also assume either model JC6, K2ST or K3ST and MC.

Corollary 3 seeks for the empirical distribution **x** of *Lophiarella* the parameter val-ues 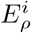 that minimize *D*(𝒯, *x*), where 𝒯 is given. The criteria for choosing (*S*_3_, *K*2*ST*) as the model explaining *Lophiarella* is MC: 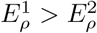 and 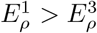 for *ρ* ∈ *𝒞* (𝒯).

We use the method of phylogenetic invariants together with the inequalities in Corollary 7 as a confirmatory conclusion that (*S*_3_, *K*2*ST*) is the model that best explains the evolution of *Lophiarella*. The phylogenetic tree of Figure 3 summarizes this conclusion.

For (*S*_3_, *K*2*ST*) as the model for *Lophiarella*, we proceeded with the maximum likelihood estimation technique and obtained the optimal parameter values 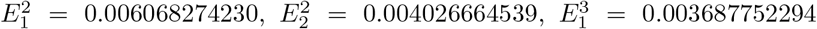 and 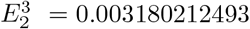, which are very similar to those obtained by the closest tree algorithm of Section 2.1.2.

### 4.1 Implications of the biological application

As shown above, we successfully inferred the phylogenetic relationships within *Lophiarella* using the the closest tree algorithm. As introduced at the beginning of this work, *Lophiarella* is a small orchid genus of the Trichocentrum clade (Orchidaceae: Oncidiinae), which is composed of three species whose distribution goes from Mexico to Nicaragua. The species are restricted to the mid to high elevations ridges and mountains. *Lophiarella splendida* and *L. microchilla* are subterrestrial or lithophilic species usually growing on exposed rocky places or cliffs of tropical or subtropical dry forest. Both species are mainly Central American and only *L. microchilla* expands its geographical distribution to Mexico, into Sierra Madre de Chiapas (600-1600 masl). Conversely, *L. flavovirens* is mainly an epiphytic species endemic to Mexico, restricted to the Mexican highlands (900-1300 masl) mainly to the western slopes of the Sierra Madre del Sur in Guerrero and the western-most slopes of the Transverse Volcanic Axis in Michoacán, Colima, and Jalisco in Tropical sub deciduous or cloud forest (Carnevali et al., 2013; Cetzal-Ix et al., 2016).

According to previous studies, the evolutionary relationships within Oncidiinae suggest that *Lophiarella* forms a monophyletic clade sister to a larger clade consisting of the genera *Cohniella, Trichocentrum* and *Lophiaris*. Within *Lophiarella*, the phylogenetic relationship inferred by (Carnevali et al., 2013) showed that *Lophiarella flavovirens* is more closely related to *L. splendida* and both species form a sister clade to *L. microchilla*. However, our results support the hypothesis that *Lophiarella microchila* is more closely related to *L. flavovirens* and both related to *L. splendida* (Figure 3). This phylogenetic relationship is consistent with the phylogenetic hypothesis based on morphological (Carnevali et al., 2013), morphological and anatomical (Cetzal Ix, 2007), and molecular evidence (Sosa et al., 2001). The topological differences could be attributed to the phylogenetic reconstruction method, since the topologies reported by (Carnevali et al., 2013) and (Cetzal Ix, 2007) are based on Parsimony, while here the topology is based on closest-tree and Hadamard methods.

The phylogenetic hypothesis found in this study (Figure 3) has a high correspondence with floral morphology since both *L. microchila* and *L. flavovirens* have small flowers and a reduced transverse labella - the differentiated median petal of the orchid flower (Dressler, 1993)-with minute triangular midlobe and a pollinarium with a long laminar stipe, which could suggest an evolutionary convergence triggered by similarities in pollinators. Furthermore, both species have a geographic correspondence, since both are distributed on the Mexican Pacific slope in tropical deciduous and subdeciduous forests. On the other hand, *L. splendida* has a very showy flower with a long, bright yellow labellum, very similar to the typical flower shape of the subtribe Oncidiinae. *L. splendida* is distributed in tropical and subtropical dry forests and, although it can occur in sympatry with *L. microchila*, the large differences in floral traits could confer pollinator specificity.

As a conclusion, the use of Phylogenetics is widespread and applicable across several biological domains, including phylogenetic inference (Duchen, 2021a,b), biogeography, and trait evolution (Duchen et al., 2017, 2020, 2021). It is precisely trait evolution what dictates morphology and it is biogeography what dictates the distribution of species. Put together, phylogenetic inference from DNA alignments is fundamental for deciphering the morphological and biogeographical evolution among species, which is exactly what we did here for *Lophiarella* by means of Hadamard conjugation and the closest-tree algorithm. Discerning the phylogenetic relationships within *Lophiarella* is not only important to understand its biogeographical and morphological evolution, but it is also relevant in conservation. For instance, two of the three species (*L. splendida* and *L. flavovirens*) are considered endangered and vulnerable, respectively, according to the IUCN. Although *L. splendida* is widely distributed (Guatemala, Honduras and Nicaragua), its showy flowers have made it one of the most popular orchids in cultivation, so its natural populations have been reduced in most areas. In contrast, *L. flavovirens* is endemic to Mexico and its populations are naturally small and distributed in discontinuous ranges, making it susceptible to extinction Carnevali et al. (2013).

## Acknowledgements

We would like to thank M.A. Steel from the University of Canterbury for his willingness to clarify issues related to the closest tree algorithm and especially for the demonstration of corollary 9, which he kindly provided us. We also thank Shu Wei Chou-chen from the University of Costa Rica for clarifying us how to perform the goodness of fit test in Section 3.2.

## Appendix

A Solution to a Special Linear System of Equations

**Lemma 8**. *Let n* ∈ ℤ^+^ *be an integer grater than* 1. *The n* × *n system of linear equations MX* = *B, where*

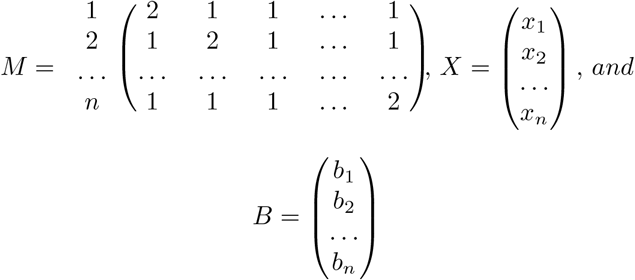

*has the unique solution*

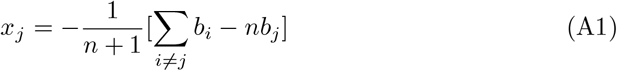

*for all j ∈ {*1, 2, 3, …, *n*}.

*Proof*. We will proceed by Cramer’s rule. For this, we will use the following notation: for each index *j* with 1≤ *j* ≤ *n, M*_*j*_ is the matrix obtained from *M*, replacing its *j*-th column by the matrix *B*. The notation *M*_*j*_(*i* |*k*) represents the minor obtained from *M*_*j*_, suppressing its *i*-th row and its *k*-th column.

To calculate the determinants Δ_*j*_ = det(*M*_*j*_), we will do Laplace expansion on the *j*-th column, which implies that we will be calculating determinants of the minors *M*_*j*_(*i*|*j*), fixing the index *j* and varying the index *i* from 1 to *n*. We also do Δ = det(*M*).

The determinant det(*M*_*j*_(*i*|*j*)) is computed case-wise on the index *i*: *i < j, i* = *j* and *i > j*.

For *i < j*, the minor *M*_*j*_(*i*|*j*) is as follows:

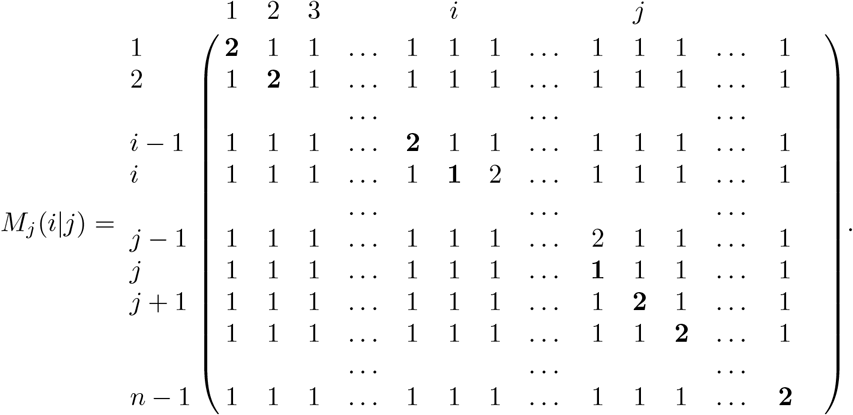

For *i* = *j*, the minor *M*_*j*_(*j*| *j*) has the same structure as *M* but of size *n−* 1 *× n −*1.For *i > j*, the minor *M*_*j*_(*i*|*j*) looks like this:

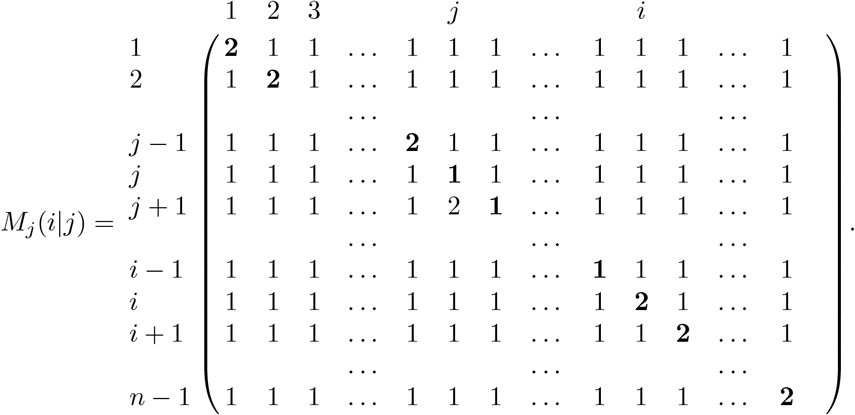

The determinant det(*M*_*j*_(*i*|*j*)) has the following formula:

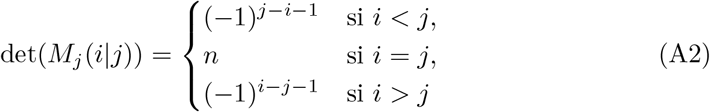

which is obtained by reducing the corresponding minor with elementary row operations. Let us reserve the symbol ∼ to denote that two matrices are equivalent (i.e. one is obtained from the other, using elementary row operations). The minor *M*_*j*_(*i*|*j*) looks as follows:

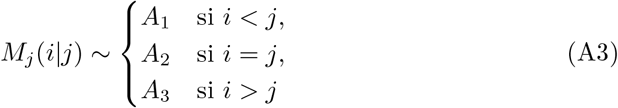

Where

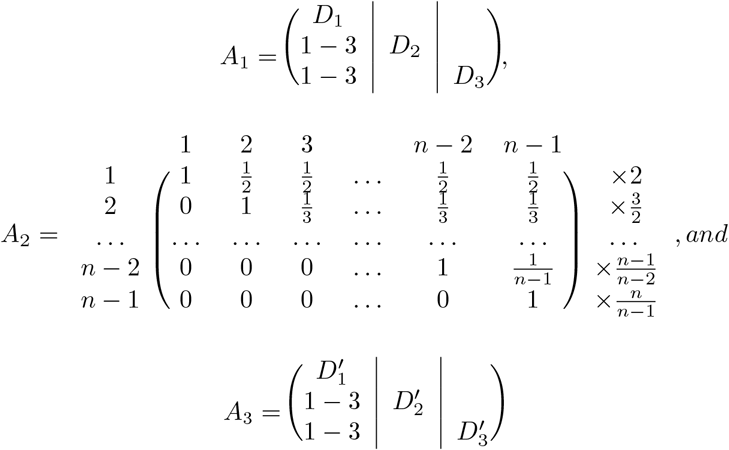

With

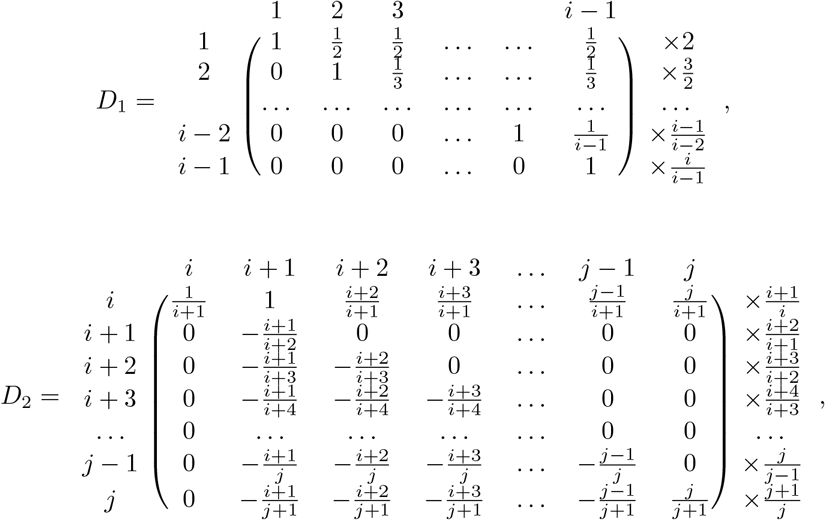

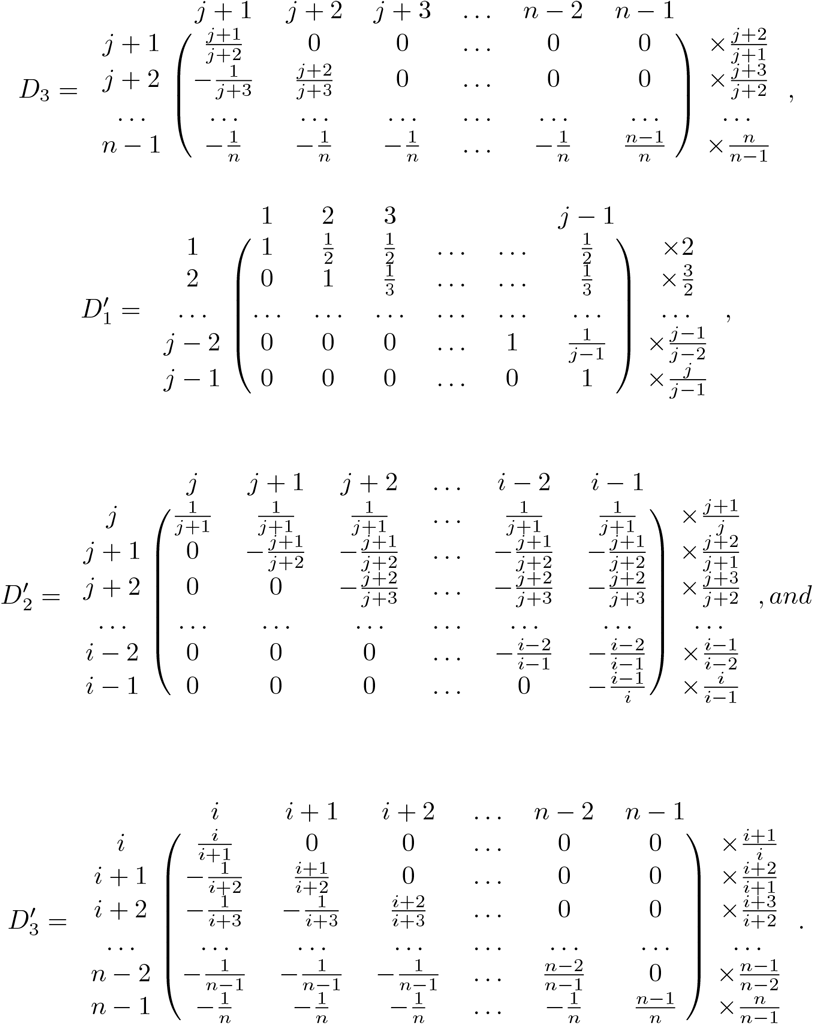

Next to the matrices 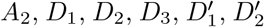 and 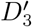 (all square matrices), we have indicated on the right the factors that compensate the calculation of the determinants. That is, for example, *M*_*j*_(*j*|*j*) ∼ *A*_2_ for all *j*, and 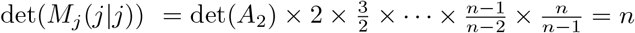, since each of the factors added to the determinant of *A*_2_ fulfills the fact that its denominator cancels the numerator of the previous factor. With this in mind, we can confirm the validity of the Formula A2.

The calculations that verify the Formula A3 are omitted, but they can be reproduced, having as a guide the intention of bringing the corresponding minor to its triangular form (upper or lower) to calculate the determinant of the equivalent matrix as a product of its terms of its main diagonal.

Let’s set an index *j* and calculate the solution 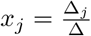 :

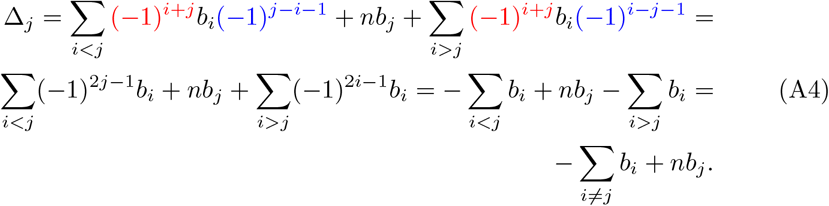

The powers of −1 highlighted in red in the Equation (A4) indicate that they come from the Laplace expansion (on the 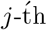 column of *M*_*j*_), while the powers of −1 highlighted in blue indicate that they come from the determinants of the corresponding minors, applying Formula A2.

Therefore

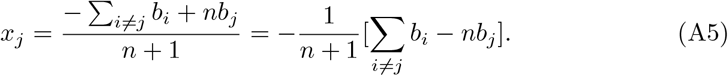

## Appendix B Approximation Formula

Corollary 3 searches for optimal values for the parameters 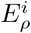 as in Section 2.1.2 when the tree topology is fixed. However, if the objective is to look for the topology of the phylogenetic tree, the following result, given by Steel et al. (1992) is preferable. The proof of Corollary 9 has been provided by Steel, M.A.

**Corolary 9**. *A closest tree for x is a tree 𝒯 that minimizes*

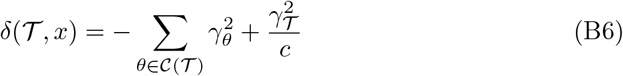

*For this tree*,

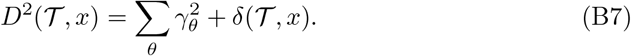

*Proof*. We proceed with Lagrange multipliers.

Let *x*_*θ*_, *θ ∈ 𝒞* (𝒯), satisfy 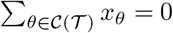.

Let *D*^2^ = Σ _*θ∈ 𝒞* (𝒯)_ (*x*_*θ*_ *− y*_*θ*_)^2^. We look for a *y* that minimizes *D*^2^ subject to conditions:

Then,

- Σ_*θ*_ *y* _*θ*_ *=0*
- Σ_*θ∉ 𝒞(𝒯)*_ *y*_θ_ = 0.

Then,

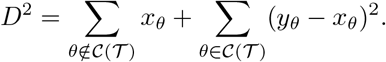

By using Lagrange multipliers, we obtain the solution

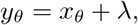

where 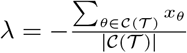.

Hence,

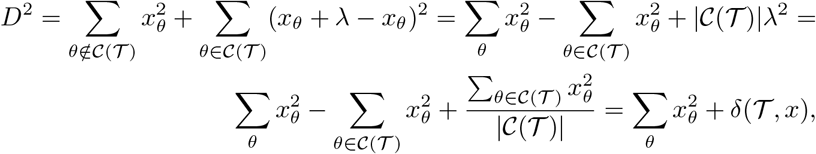

where 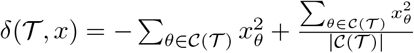 and 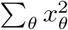 is constant.

